# When batch correction corrupts gene expression: uncovering distortions in correlation structures

**DOI:** 10.64898/2026.06.06.729466

**Authors:** Jalil Nourisa, Antoine Passemiers, Yves Moreau, Daniele Raimondi

## Abstract

Batch correction is essential for integrating datasets and enabling population-level insights into health and disease. Embedding-based approaches are among the most widely used solutions, but here we highlight a critical, overlooked limitation: these methods can distort feature-to-feature (e.g., gene–gene) relationships, potentially undermining downstream analyses. We investigate this issue and introduce a novel metric to quantify it.

## 1 Introduction

Batch correction is a crucial step in processing single-cell omics to address unwanted technical variability that arises from different experimental batches [1]. This variability, often introduced by differences in sample preparation, sequencing runs, or reagent batches, can not only obscure true biological signals (making it challenging to compare data between batches), but also lead to misleading biological interpretations and spurious conclusions. Batch correction methods aim at alleviating these effects by disentangling the biological differences from the technical artifacts, while both contribute to the overall data variance [2].

Numerous algorithms have been developed for single-cell data integration, offering corrections at various levels: feature-level [3, 4, 5, 6], embedding-level [7, 8], and graph-level [7]. Feature-level integration generates a corrected data matrix, such as gene expression, enabling downstream analyses like differential expression, gene correlation studies, and perturbational modeling. These methods employ diverse approaches to address batch effects. For instance, global methods like ComBat [3] model batch effects as additive or multiplicative shifts affecting all cells uniformly. On the other hand, embedding-level methods project features into a lower-dimensional space, align cells within this space to remove batch effect, and optionally map the data back to the original feature space. These approaches can utilize linear techniques, such as Seurat and Scanorama [9, 4], or deep learning frameworks, particularly (variational) autoencoders, as implemented in scVI, scGen, and scALEX [8, 5, 6].

An ideal batch correction method would eliminate technical effects without distorting the underlying biological data. However, achieving this balance is challenging, especially in datasets with more complex batch effects [2]. In the absence of a universally optimal solution, researchers often aim for a ‘silver standard’ approach that effectively mitigates batch effects while minimizing the introduced bias. Independent benchmark studies have played a pivotal role in this pursuit, standardizing performance evaluations across diverse datasets [10, 2]. For instance, Luecken et al. [2] introduced 14 metrics to evaluate both batch effect removal and the preservation of biological signals. Despite these advances, the assessment of biological conservation remains relatively underexplored [11, 12]. In particular, unsupervised metrics proved to be insufficient for detecting residual batch effects, especially when the underlying relationship is non-linear [12].

While embedding-based batch correction methods have become standard practice in single-cell data integration, their effects on preserving biologically meaningful gene-gene relationships remain largely unexplored. In this study, we take a closer look at how common integration steps—particularly dimensionality reduction and reconstruction—can distort gene-gene correlation structures that are critical for downstream analyses. We show that both linear and non-linear embeddings can introduce substantial redundancy into the data, ultimately impairing downstream analyses such as gene regulatory network (GRN) inference. To address this, we introduce a new metric based on feature-feature relationship (FFR) conservation, that directly quantifies the preservation of biologically relevant feature relationships. Our results highlight the need for careful consideration of integration strategies, especially when the preservation of gene-level relationships is essential for biological interpretation.

## 2 Results

### 2.1 Impact of embedding-based integration on gene correlation structure

Embedding-based batch correction approaches typically follow a paradigm of data embedding, batch alignment, and data reconstruction. To demonstrate how this process introduces distortions into FFRs, we first examined the effects of data embedding and reconstruction (auto-encoding) on gene correlations. Specifically, we embedded single-cell gene expression data into a low-dimensional space using principal component analysis (PCA) with varying numbers of principal components (PCs) and then reconstructed the data, without any correction (Methods). Pair-wise Pearson correlations were calculated between genes for both the original and reconstructed datasets to evaluate how reconstruction affects these correlations.

Dimensionality reduction with smaller numbers of PCs (e.g., 5, 10, and 100) significantly inflated gene-gene correlations compared to the original data (Fig. 1-A). Additionally, analysis of the top 20 gene-gene links with the highest correlations in the original dataset revealed substantial alterations in correlation rankings after reconstruction. Some links that were among the original top 20 dropped dramatically in importance, with a few falling to rankings beyond 10,000—a threshold we capped for visualization (Fig. 1-A). It must be noted that these significant distortions have been introduced by mere principal component analysis. In practice, sophisticated embedding methods (such as VAE, which are more common in the single-cell literature) have even more room for performing arbitrary data modifications due to their non-linear nature. Therefore, we conducted the same analysis on batch correction methods to further evaluate this issue.

**Figure 1:**
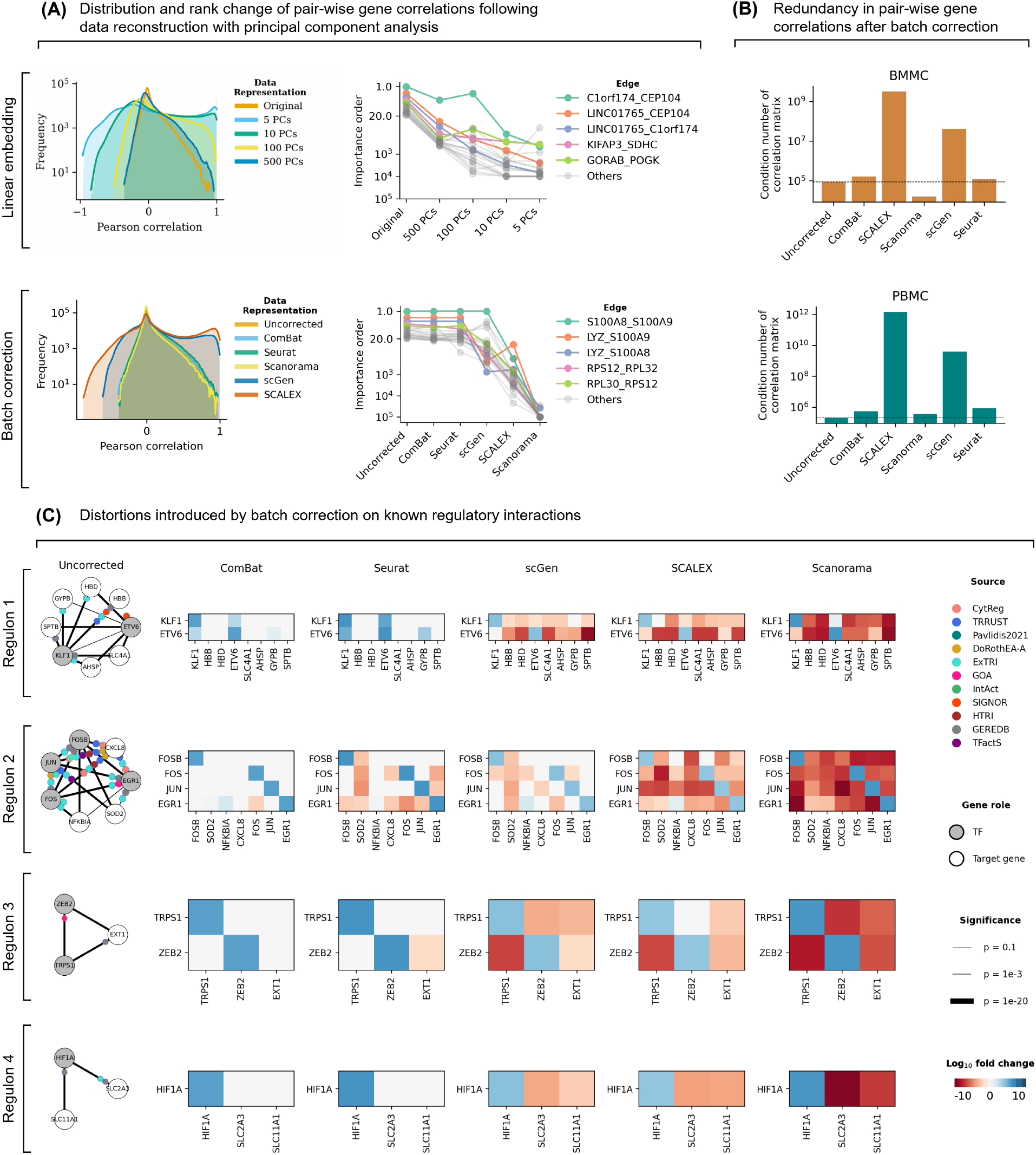
(A) Evaluation of pairwise gene correlations in original and reconstructed data following dimensionality reduction (first batch of the BMMC dataset) and batch correction (entire PBMC dataset). Upper row assesses the effect of linear embedding using varying numbers of principal components (PCs), while bottom row examines different batch correction methods. The left column shows the distribution of pair-wise Pearson correlations between genes. The right column displays the change in the ranking of the top 20 gene-gene correlations (edges) based on the original data, highlighting how embedding or correction methods impact the strongest interactions. (B) Redundancy in the gene correlation matrix before and after batch correction. We quantify informational redundancy in the pairwise gene correlation matrix using the condition number—a continuous measure of collinearity. Results are shown for the BMMC and PBMC datasets. (C) Evaluation of known regulons after batch correction. Regulons were inferred for myeloid cells using GENIE3 on the BMMC dataset after applying different batch correction methods. Link confidence is encoded by edge thickness. Each CollecTRI source is represented by a uniquely colored circle. TF regulators are shown in grey. Log fold changes in edge *p*-values are displayed as heatmaps.

We extended our analysis to real batch correction scenarios using commonly employed methods, including ComBat, Seurat [4], Scanorama [9], scGen [5], and SCALEX [6] (Methods). ComBat uses a global approach in correcting feature space and therefore does not rely on embeddings. Seurat employs a linear embedding approach with mutual nearest neighbors (MNN) to align datasets. Scanorama utilizes a non-linear embedding approach with manifold alignment to integrate datasets and correct batch effects. Both scGen and SCALEX adopt a variational auto-encoder (VAE) architecture for dimensionality reduction and integration. For the latter four approaches, we anticipate varying degrees of distortions.

By evaluating pair-wise Pearson correlations between genes, our results showed that scGen and SCALEX notably inflated correlation values compared to the uncorrected data (Fig. 1-A). Examining the top 20 gene-gene correlations revealed that, while ComBat and Seurat introduced minimal alterations, Scanorama, scGen, and SCALEX substantially distorted the order (rank) of these top edges.

#### 2.2 Dimensionality reduction increases redundancy in feature-feature correlation structures

To empirically assess the extent of the distortion introduced by both linear and non-linear embeddings, we used two methods. First, we derived a mathematical formalism to describe how dimensionality reduction through linear embedding affects the feature-feature correlation matrix (FFR), showing that this process increases collinearity and can distort the underlying correlation structure (see Methods). Second, we employed the condition number—a continuous metric closely related to matrix rank—to quantify the degree of redundancy introduced into the correlation matrix during embedding. Across datasets, we found that all batch correction methods increased the condition number of the correlation matrix, with the sole exception of Scanorama on the BMMC dataset (Fig.1-B). While linear embedding methods such as Seurat and Scanorama might be expected to exhibit higher collinearity, it was in fact the VAE-based methods (SCALEX and scGen), which use non-linear embeddings, that resulted in the largest condition numbers. These findings indicate that embedding-based methods can introduce substantial redundancy into the reconstructed feature space, potentially limiting the interpretability and robustness of downstream correlation-based analyses.

#### 2.3 Batch correction impairs gene regulatory network inference

GRN inference aims to reconstruct the regulatory relationships between transcription factors (TFs) and their target genes from gene expression or multimodal data. Considering that transcriptomics-based GRN inference primarily relies on gene-gene covariance or co-expression patterns, we hypothesized that batch correction—which we have shown to introduce unwanted biases into FFRs—could also impair the accuracy of GRN reconstruction. To test this hypothesis, we inferred GRNs for five major immune cell populations: CD4^+^ T cells, CD8^+^ T cells, myeloid cells, natural killer (NK) cells, and B cells. For each cell type, we applied three established GRN inference algorithms: GENIE3 [13], ENNET [14], PORTIA [15]. GRNs were constructed using the uncorrected datasets, as well as the same datasets preprocessed with the batch correction methods previously evaluated.

To quantify the impact of batch correction on the biological fidelity of inferred GRNs, we performed two complementary analyses (see Methods). First, we inferred GRNs across all combinations of dataset, cell type, batch correction method, and inference algorithm. Each GRN was evaluated against the CollecTRI database [16], which contains experimentally validated TF–target interactions and serves as a proxy ground truth. For each GRN, we computed the area under the receiver operating characteristic curve (AUROC) to assess how well the inferred TF–gene edges aligned with ground-truth interactions. To evaluate the significance of performance changes between the uncorrected dataset and its corrected versions, we applied an ordinary least squares (OLS) regression model [17], which allowed us to disentangle the contributions of dataset, cell type, batch correction method, and inference algorithm to the AUROC scores. The contribution and significance of each factor are reported in Supplementary Table S1. We found that all batch correction methods reduced AUROC scores (Supplementary Figure 1), with significant decreases observed for Seurat (−0.0086, *p <* 0.005) and Scanorama (−0.0150, *p <* 0.005) (Supplementary Table S1).

To further evaluate the quality of GRN reconstruction in more details, we evaluated TF-target pairs centered around key TFs with well-established roles. To this end, we focused on the GENIE3-inferred GRNs for myeloid cells, which achieved the highest AUROC score among all GRN models (Table S1). We selected 4 subgraphs based on the consensus between CollecTRI and GENIE3 (Methods). These subgraphs include TFs such as *FOS, FOSB*, and *JUN* —subunits of the AP-1 TF complex—with known roles in myeloid cell differentiation and inflammatory responses [18, 19]. For both the uncorrected GRN and each batch-corrected version, we transformed regulatory weights into *p*-values under a normality assumption. Next, we computed the log fold changes in *p*-values between the uncorrected and batch-corrected GRNs. Our analysis revealed that GRNs inferred from uncorrected data consistently retained statistically significant edges supported by CollecTRI. In contrast, most batch correction methods—except ComBat—frequently reduced the statistical significance of these edges (as shown by the red grid cells in the heatmaps; Fig. 1-C). This effect was particularly pronounced for Scanorama, scGen, and SCALEX. For example, *JUN–CXCL8* and *FOS–CXCL8* edges were strongly significant (*p <* 10^*−*10^) in the uncorrected data and supported by multiple sources in Collec-TRI, yet these links were completely lost in the GRNs inferred from Scanorama- and SCALEX-corrected data. A similar pattern was observed for *KLF1–HBB*. Additionally, scGen, SCALEX, and Scanorama all negatively affected the detection of *HIF1A–SLC11A1* and *HIF1A–SLC2A3* interactions, while *HIF1A* has a well-known role in mediating hypoxia responses in myeloid cells during inflammation and infection [20]. These results further support that embedding-based batch correction methods can substantially distort biologically meaningful regulatory interactions in GRNs.

#### 2.4 A new metric to assess batch correction quality: FFR conservation

Most of the biological conservation metrics listed in [2] are based on clustering and how clusters relate to cell types. Here, we designed a more straightforward and intuitive metric, termed the *FFR conservation*, to evaluate the impact of batch correction on FFRs (Methods). This metric assesses how well the relationships between features (e.g., genes) are preserved following batch correction. Specifically, the *FFR conservation* focuses on the most significant FFRs, calculating both the recall of these important edges post-correction (*FFR recall*) and the Spearman correlation between features pre- and post-correction (*FFR order*).

Using the *FFR conservation*, we evaluated the quality of batch correction for five methods across two datasets (Methods). Additionally, we compared the correction quality using 13 established metrics from scBI [2], as well as the BPS-RF score from [12] (Methods). The results showed that ComBat achieved near-perfect performance in terms of *FFR conservation*, followed by Seurat (Fig. 2). In contrast, methods relying on dimensionality reduction—scGen, SCALEX, and Scanorama—significantly compromised FFR preservation. Notably, none of the existing metrics for biological conservation reflected a trend similar to what is observed for *FFR conservation*, highlighting the unique aspects of biological signal retention captured by this metric.

**Figure 2:**
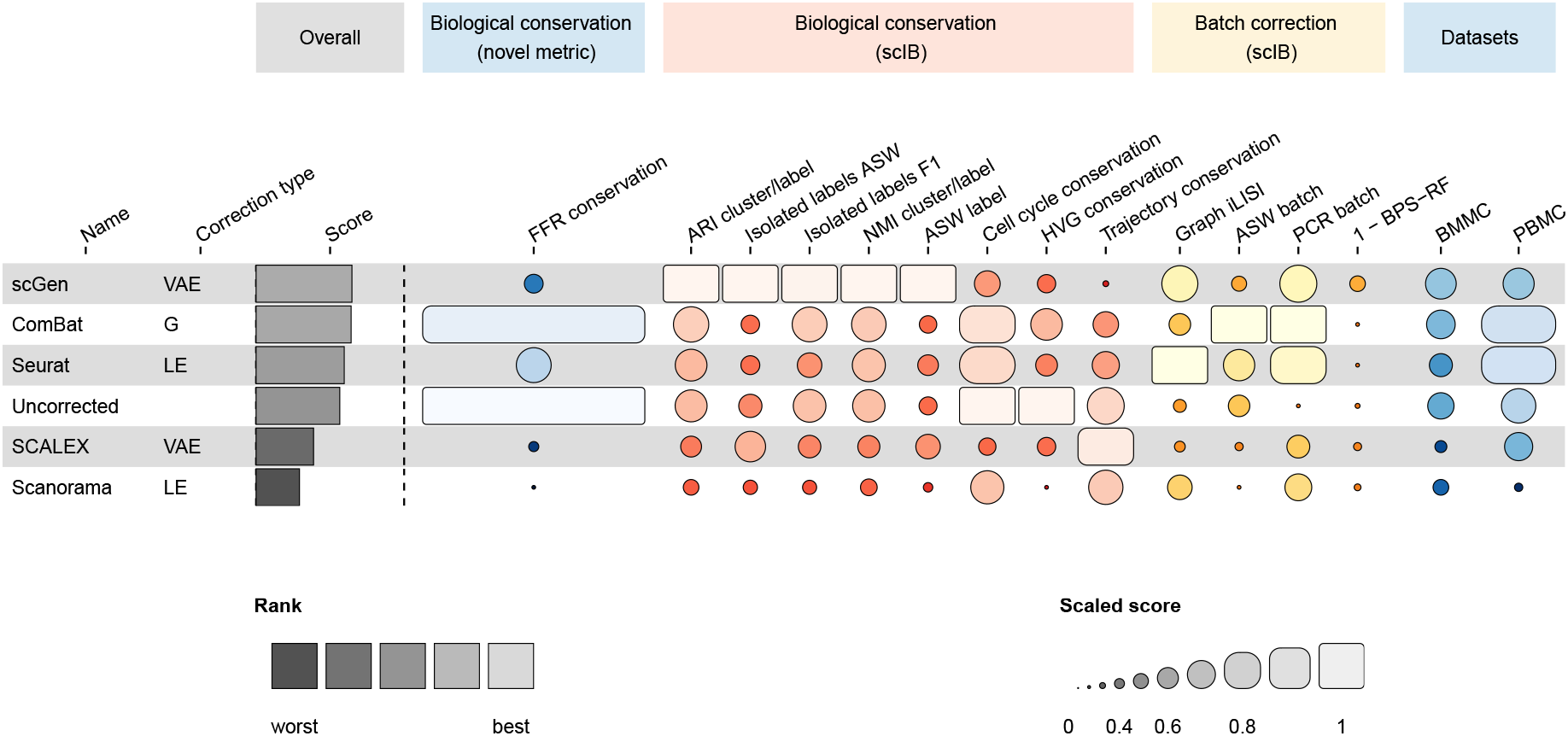
Summary of batch correction performance. The different methods are annotated with their underlying correction approach; variational auto-encoder (VAE), linear-embedding (LE), and global (G). The correction efficiency was evaluated across two datasets: peripheral blood mononuclear cells (PBMC) and bone marrow mononuclear cells (BMMC). Of note, we excluded two metrics—Graph cLISI and Graph Connectivity—as they exhibited minimal variation across methods.

Finally, all batch correction methods produced a BPS-RF strictly higher than 0.6 and close to 1 for most methods (ComBat and Seurat even increased the BPS-RF compared w.r.t. the uncorrected data), which demonstrates their inability to fully remove residual batch signals.

## 3 Conclusion

In this work, we demonstrated that feature-level batch correction methods based on embeddings—particularly those employing dimensionality reduction—can substantially distort feature–feature relationships (FFRs). Such distortions include the removal of authentic FFRs and the introduction of redundant correlations, indicating significant information loss. We exemplified this issue in the context of GRN inference, where batch correction was shown to reduce the efficiency of network reconstruction and to eliminate well-established regulatory connections. These distortions are likely to compromise other downstream analyses that rely on accurate feature representations, such as co-expression analysis, co-accessibility analysis, or perturbation response prediction. To address this issue, we introduced a novel metric that quantifies the extent of correlation-structure distortion and provides guidance for selecting appropriate batch correction methods.

We emphasize the need for systematic evaluation of correction strategies, with particular attention to their ability to preserve biological signal and minimize bias. Such evaluations should incorporate task-specific performance metrics to ensure that correction methods remain aligned with the objectives of downstream analyses.

## 4 Data availability statement

The two datasets used in this manuscript were obtained from https://www.ncbi.nlm.nih.gov/geo/query/acc.cgi?acc=GSE1941 and https://www.ncbi.nlm.nih.gov/geo/ respectively.

## 5 Methods

### 5.1 Datasets

Two datasets were used in this report: the BMMC dataset, taken from the Open Problems in Single-Cell Analysis NeurIPS Competition 2021 [21], the PBMC dataset obtained from [22]. We took five batches from each dataset. Both datasets included single-cell RNA sequencing (scRNA-seq) data, preprocessed to exclude low-quality cells (minimum of 10 detected genes) and genes (expressed in at least 100 cells). They were cell-type annotated and contained between 5,000 and 8,000 single cells per batch. Normalization was performed using the shifted logarithmic approach (SLA) via *Scanpy*, applying *normalize total* followed by *log1p* transformation [23]. We refer to the resulting data as the uncorrected dataset.

### 5.2 Linear embedding using principal component analysis

Expression data was embedded it into lower-dimensional spaces using 5, 10, 100, and 500 components with singular value decomposition (SVD) from *scikit-learn*. The data was constructed back to the original feature space using inverse embedding for each number of dimensions. Next, for the uncorrected data and each of the reconstructed dataset, we calculated the Pearson correlations between genes. It is important to note that here we do not correct the batch effects. Also, we only used data from one batch of BMMC (with single cell count of over 5000) to avoid batch effects.

### 5.3 Calculation of the *FFR conservation*

We select *b* individual batches (in this report, *b* = 5) with sufficient single cells (at least 5,000 per batch) to reliably infer correlations. For each batch *i*, we calculate the feature correlation matrix **C**^(*i*)^ *∈* ℝ^*m×m*^ using the uncorrected data independently, where *i ∈* {1, …, *b*}, and *m* represents the number of features (genes), ensuring no batch effects are introduced. To reduce data dimensionality, we select the 1,000 most highly variable genes. Similarly, we compute the correlations for the same selected links in the corrected data, denoted as 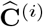.

The *FFR recall* (*S*_recall_) quantifies the overlap between the top *q* links (set to *q* = 1, 000) in the uncorrected (**C**_(i)_) and corrected 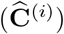 datasets. Let 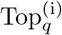 and 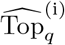 denote the sets of the top *q* feature-to-feature links for the uncorrected and corrected *i*-th batch, respectively. The FFR recall is defined as:

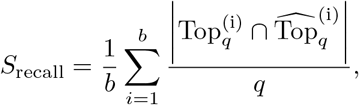

where *S*_recall_ *∈* [0, 1], with higher values indicating better overlap between the uncorrected and corrected top links.

To compute the *FFR order* (*S*_order_), we take the top *q* edges from the uncorrected correlation matrix (**C**^(*i*)^) and calculate the Spearman correlation *ρ* between their ranks in **C**^(*i*)^ and 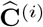. The FFR order is given by

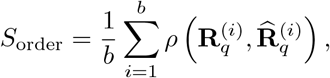

where 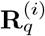 and 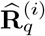 denote the ranks (importance order) of the top *q* edges in the uncorrected and corrected datasets, respectively. Higher *S*_order_ values indicate better preservation of the relative importance of the top edges.

Finally, the *FFR conservation* score (*S*_FFR_) is computed as the average of these two metrics:

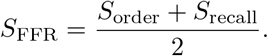

This score provides a comprehensive measure of the preservation of feature-to-feature relationships, with higher values indicating higher performance.

### 5.4 Batch correction evaluation metrics

We took 13 metrics from scBI [2], as well as the BPS-RF score defined in [12]. We provide a summary of these metrics here. The metrics measuring batch resolving are

- **Graph iLISI:** Integration Local Inverse Simpson Index (iLISI) quantifies the mixing of batches in the corrected data, with higher values indicating better batch mixing.
- **ASW Batch:** Average silhouette width (ASW) evaluates how well batch-specific clustering is removed in the corrected data.
- **PCR Batch:** The Principal Component Regression (PCR)-based score assesses the extent to which batch effects are removed from the principal components of the data.
- **BPS-RF:** The metric is defined as 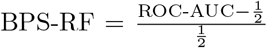, where ROC-AUC is the area under the ROC curve obtained through stratified 5-fold cross-validation of a random forest classifier predicting the batch from the gene expression data directly. We used the same hyper-parameters as in the original study [12].

The metrics measuring biological conservation are:

- **ARI Cluster/Label:** The adjusted Rand Index (ARI) evaluates how well the Leiden clusters correspond to cell type annotation.
- **Isolated Labels ASW:** ASW quantifies how well isolated cell populations (labels) are separated in the corrected data.
- **Isolated Labels F1:** The F1-score assesses the classification performance for isolated cell populations, balancing precision and recall.
- **NMI Cluster/Label:** Normalized Mutual Information (NMI) measures the similarity between clustering results and ground truth labels, accounting for entropy.
- **ASW Label:** ASW for cell type labels evaluates the separation of clusters based on known cell types.
- **Cell Cycle Conservation:** Assesses how well the corrected data retains biological variability associated with the cell cycle.
- **HVG Conservation:** Evaluates the preservation of highly variable genes (HVGs) after batch correction.
- **Trajectory Conservation:** Measures how well developmental trajectories or pseudotime orders are preserved in the corrected data.

### 5.5 Gene regulation network inference and regulon selection

#### 5.5.1 GRN inference

Using transcriptomic data from two datasets—PBMC and BMMC—we inferred GRNs for five major immune cell types: CD4^+^ T cells, CD8^+^ T cells, myeloid cells, NK cells, and B cells using three established methods: GENIE3 [13], PORTIA [15], and ENNET [14]. PORTIA was run with default hyperparameters, except the Box-Cox transformation was disabled using `method=‘no-transform’` to accommodate non-strictly positive input values. ENNET and GENIE3 were both run with default settings. Each method was supplied with the predefined list of TFs as candidate regulators. As a baseline, we also constructed a random network consisting of a TF–target interaction matrix where each entry was sampled independently from a uniform distribution over [0, 1]. GRN inference was repeated for each version of the datasets: uncorrected, and batch-corrected using scGen, SCALEX, Scanorama, Combat, and Seurat. Let’s note that ENNET failed to infer 10 out of the 60 networks.

#### 5.5.2 Assessing the effect of batch correction on GRN fidelity

We evaluated the accuracy of the inferred GRNs using CollecTRI as a proxy gold standard (GT) [24, 25], which provides curated TF–target gene interactions. To map the inferred GRNs to the GT, we retained only those TFs from the inferred networks that regulate at least one target gene in CollecTRI. Similarly, we included all CollecTRI target genes that were present in the dataset. Then, we computed the area under the receiver operating characteristic curve (AUROC) between the inferred interactions and the ground truth. This evaluation was performed 240 times in total: across 2 datasets, 6 preprocessing versions, 5 cell type categories, and 4 GRN inference methods.

Next, we fitted an ordinary least squares (OLS) regression model using the `statsmodels` Python package [17], using the AUROC as response variable, and using the dataset, version, cell type category and GRN inference method as explanatory categorical variables. The coefficients of such a model represent the linear contributions of these latter variables to the performance, and the corresponding *t*-test *p*-values reflect the significance of these contributions. “Uncorrected” and “random” were selected as the default categories for the dataset version and the GRN inference method, respectively. These categories are therefore absent from the first column of Table S1. In such a way, the linear coefficients of ComBat, SCALEX, Seurat, scGen and Scanorama reflect their relative improvement (i.e., contrast) over the absence of batch correction. Analogously, the coefficients of PORTIA, GENIE3 and ENNET should be interpreted as the relative improvement of each of these methods over a purely random GRN.

#### 5.5.3 Identifying regulons of interest

In order to evaluate to which extent batch correction preserves well-understood inferred regulatory links, we focused on the inferred GRN of highest reliability, namely the myeloid cell GRN inferred by GENIE3 from the uncorrected BMMC dataset (AUROC=0.577). We first sorted the TF–target edges by decreasing order of confidence, and filtered out the ones not present in the CollecTRI. Next, we selected the top 1,000 ones, and extracted subgraphs from the resulting edge list. Each subgraph was constructed iteratively by identifying genes connected to one of its genes via one of the edges in that list. We considered only subgraphs of size *≥* 3, which resulted in 4 subgraphs.

### 5.6 Mathematical formulation of linear embedding-based data transformation

The correlation matrix distortion can be formalized as:

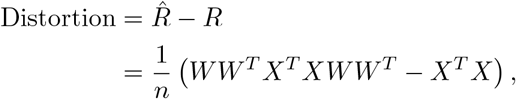

where *R* and 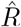 are the correlation matrices before and after batch correction, respectively. *X* is the centered gene expression and *W* is the linear projection corresponding to the principal component analysis of *X*.

This metric captures the extent to which the gene-gene correlation structure is distorted. Additionally, we employed the condition number of 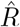 to quantify the degree of redundancy present in the correlation matrix. The condition number is defined as:

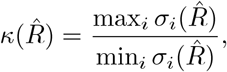

where 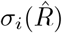 is the *i*th singular value of 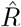.

More mathematical details are provided in Supplementary Section 1.

