## Supplementary data for "When batch correction corrupts gene expression: uncovering distortions in correlation structures"

### Supplementary materials for the manuscript "When batch correction corrupts gene expression: uncovering distortions in correlation structures"

June 2, 2026

#### 1 Mathematical formulation of linear embedding-based data transformation

Here, we examine the process of data transformation during PCA embedding. For a data matrix $X \in \mathbb{R}^{n \times m}$, where $n$ is the number of samples and $m$ the number of features (e.g., genes), PCA projects the data onto a lower-dimensional space defined by $p$ ($\leq m$) principal components. This projection yields a component matrix $Z \in \mathbb{R}^{n \times p}$ such that:

$$Z = XW,$$

where $Z_{ij}$ is the $i$-th sample's coordinate on the $j$-th principal component, and $W_{kj}$ is the loading of feature $k$ on component $j$. Matrix $X$ is assumed to be centered and scaled.

To approximately reconstruct $X$ from $Z$, we use the inverse transformation:

$$\hat{X} = ZW^T.$$

Expanding $Z$ in terms of $X$ from the previous step gives:

$$\hat{X} = XWW^T.$$

Here, $WW^T$ acts as a coupling matrix during reconstruction, effectively blending information across features. Let's note that $W$ is itself a non-linear function of $X$, as it is generally obtained through the singular value decomposition of $X$. This cross-feature influence in the reconstructed $\hat{X}$ matrix is what introduces the distortion: each feature (column) in $\hat{X}$ is no longer solely reflective of the original column in $X$, but also contains contributions from other features, creating artificial correlations.

To quantify the bias added by PCA into FFRs, we express feature-feature correlations as a matrix $\hat{R}$:

$$\hat{R} = \frac{1}{n}\hat{X}^T\hat{X},$$

where $\hat{R}_{ij}$ is equal to the correlation between reconstructed features $i$ and $j$. This holds when the original features are first centered and scaled.

Substituting $\hat{X}$ in the previous formula gives:

$$\hat{R} = \frac{1}{n} W W^T X^T X W W^T = W W^T R W W^T.$$

Finally, the distortion of the correlation matrix can be formalized as:

$$\begin{aligned}\text{Distortion} &= \hat{R} - R \\ &= \frac{1}{n}\left(W W^T X^T X W W^T - X^T X\right)\end{aligned}$$

Assuming $X$ is full-rank, $R$ is a $m \times m$ symmetric positive definite matrix with rank $m$, and the rank of matrix $W$ is $p \leq m$. Therefore, the rank of matrix $\hat{R}$ is equal to the minimum of ranks, hence $p$. This results in a rank loss of $m - p$ and therefore an information loss regarding the correlation matrix. In terms of degrees of freedom, a rank-$p$ symmetric matrix of dimensions $m \times m$ has $p(2m - p)$ degrees of freedom. Therefore, PCA introduces a reduction of $m^2 - p(2m - p)$ degrees of freedom, which can be attributed to an information bottleneck. Importantly, this constraint is a non-linear function of the original matrix (since $WW^T$ depends itself on $X$ through SVD), distorting the true biological relationships captured in the data.

In practice, all correlation matrices are full-rank due to the presence of noise, making matrix rank an unreliable indicator of redundancy. Instead, we employ the condition number of the reconstructed correlation matrix $\hat{R}$ to assess collinearity among features. The condition number is defined as:

$$\kappa(\hat{R}) = \frac{\max_i \sigma_i(\hat{R})}{\min_i \sigma_i(\hat{R})},$$

where $\sigma_i(\hat{R})$ is the $i$th singular value of $\hat{R}$.

This metric captures the extent to which features in $\hat{R}$ are linearly dependent. A low condition number suggests that the features are relatively independent, while a high condition number indicates that many features are nearly linear combinations of each other—signaling excessive collinearity. Since dimensionality reduction techniques reduce the intrinsic dimensionality of the data, the reconstructed features in $\hat{X}$ may inherit artificial dependencies. As a result, $\hat{R}$ becomes increasingly ill-conditioned when more information is lost (i.e., as $p \ll m$). The condition number thus serves as a proxy for the bias introduced by the embedding, quantifying how much the recovered correlation structure deviates from the true, high-dimensional relationships.

### 2 GRN inference

#### 2.1 Performance of each batch correction method

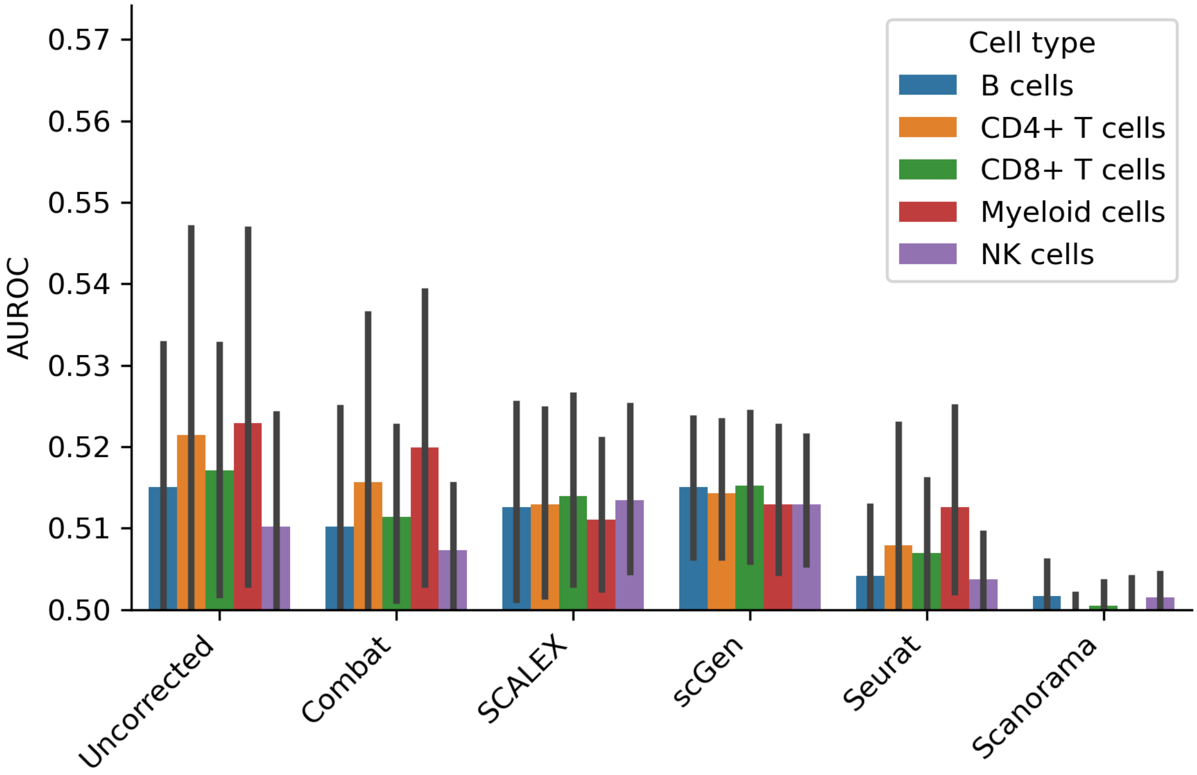


Figure 1: AUROC score of inferred GRNs for each cell type and batch correction method. Error bars represent the 95% confidence interval across the remaining variables (dataset and GRN inference method).

#### 2.2 Significance of GRN inference performance changes introduced by batch correction

| | | | | | Confidence interval | |
|---|---|---|---|---|---|---|
| Variable | Coefficient | Std. err. | $t$ | $p$-value | 0.025 | 0.975 |
| (Intercept) | 0.2972 | 0.001 | 211.860 | 0.000 | 0.294 | 0.300 |
| ComBat | -0.0026 | 0.003 | -0.941 | 0.348 | -0.008 | 0.003 |
| SCALEX | -0.0027 | 0.003 | -0.973 | 0.331 | -0.008 | 0.003 |
| Scanorama | -0.0150 | 0.003 | -5.412 | 0.000 | -0.020 | -0.010 |
| scGen | -0.0014 | 0.003 | -0.517 | 0.606 | -0.007 | 0.004 |
| Seurat | -0.0086 | 0.003 | -3.103 | 0.002 | -0.014 | -0.003 |
| ENNET | 0.0036 | 0.002 | 1.645 | 0.101 | -0.001 | 0.008 |
| GENIE3 | 0.0312 | 0.002 | 14.919 | 0.000 | 0.027 | 0.035 |
| PORTIA | 0.0060 | 0.002 | 2.864 | 0.005 | 0.002 | 0.010 |
| B cells | 0.0585 | 0.002 | 38.059 | 0.000 | 0.055 | 0.062 |
| CD4+ T cells | 0.0604 | 0.002 | 39.279 | 0.000 | 0.057 | 0.063 |
| CD8+ T cells | 0.0595 | 0.002 | 38.724 | 0.000 | 0.056 | 0.063 |
| Myeloid cells | 0.0620 | 0.002 | 40.805 | 0.000 | 0.059 | 0.065 |
| NK cells | 0.0569 | 0.002 | 36.713 | 0.000 | 0.054 | 0.060 |
| BMMC | 0.1495 | 0.001 | 145.761 | 0.000 | 0.147 | 0.152 |
| PBMC | 0.1477 | 0.001 | 142.667 | 0.000 | 0.146 | 0.150 |

Table S1: Linear contribution of each variable (dataset, batch correction method, GRN inference method, and cell type) to the AUROC score, based on an OLS regression model. The associated $p$-values were derived from $t$-tests on the model coefficients. For the batch correction factor, comparisons were made between corrected and uncorrected data. For the GRN inference method, comparisons were made between the inferred GRNs and randomly generated networks (Methods)
